# Likelihood-based Fits of Folding Transitions (LiFFT) for Biomolecule Mapping Data

**DOI:** 10.1101/294041

**Authors:** Rhiju Das

## Abstract

**Summary:** Biomolecules shift their structures as a function of temperature and concentrations of protons, ions, small molecules, proteins, and nucleic acids. These transitions impact or underlie biological function and are being monitored at increasingly high throughput. For example, folding transitions for large collections of RNAs can now be monitored at single residue resolution by chemical mapping techniques. LIkelihood-based Fits of Folding Transitions (LIFFT) quantifies these data through well-defined thermodynamic models. LIFFT implements a Bayesian framework that takes into account data at all measured residues and enables visual assessment of modeling uncertainties that can be overlooked in least-squares fits. The framework is appropriate for multimodal techniques ranging from chemical mapping including multi-wavelength spectroscopy.

**Availability:** Freely available MATLAB package at https://ribokit.stanford.edu/LIFFT/.

**Supplementary information:** Supplementary data are available at *Bioinformatics* online.

---

Biomolecules and their complexes underlie all core processes in molecular biology and human disease and their functions typically depend on a panoply of folded, partially folded, and unfolded states. Probing these states is a major part of modern biochemistry and biophysics and is the focus of important technology development. For example, RNA chemical mapping, also known as structure mapping or footprinting, can measure the reactivity of each residue of an RNA to one or more chemical modification reagents [(Kwok *et al.,* 2015) and refs. therein]. Mapping these chemical reactivities as a function of the biological partners’ concentrations or of solution conditions provides a route to discover and quantitatively characterize structural transitions of RNA.

After acquiring rich, multi-residue data for a structural transition, the next step is to fit and summarize these data in terms of a defined thermodynamic model. However, it is often difficult to assess uncertainties in these models. To address this problem for RNA chemical mapping, we developed an in-house Bayesian tool to scan over model parameters and to visualize possible correlations or nonlinearities in their uncertainties. The tool has been effective in numerous systems, including noncanonical RNA motifs stabilized through computational design (Das *et al.,* 2010), glycine-sensing bacterial riboswitches (Kladwang *et al.,* 2012), a metal ion core in a group I self-splicing intron (Frederiksen *et al.,* 2012), RNA structures designed through the internet-scale design project Eterna (Lee *et al* 2014; Cordero and Das, 2015), and chemically modified RNAs interacting with the Puf protein (Vaidyanathan *et al.,* 2017). The resulting software has now been named LIkelihood-based Fits of Folding Transitions (LIFFT) and is being made available for wider use, including applications beyond RNA mapping.

Given experimental data *D*_*ij*_ measured for a molecule under measurement conditions *i* = 1,2,…*N* (e.g., different Mg^2+^ concentrations) at residues (or wavelengths) *j =* 1, 2,… *M*, the goal is to fit this matrix of data to a model 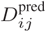:

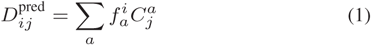

where 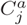 are the predicted data observable for each assumed state *a* (e.g., unfolded and folded) and 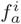 are the fractions of the molecule population in each state *a* under each condition i, which depend on the assumed thermodynamic model. A common use case involves fitting Mg^2+^-dependent RNA folding to a Hill binding curve:

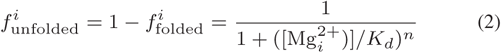

where the two fit parameters are the apparent dissociation constant *K*_*d*_ and apparent Hill coefficient *n* (Lipfert *et al.,* 2014). Bayes’ theorem gives the appropriate probability for assessing such a model. The posterior probability of the model given the data is:

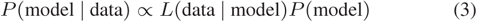

The likelihood *L*(data | model) is assumed to take a Gaussian form involving (initially unknown) experimental errors *σ*_j_ at each residue (or wavelength) *j*:

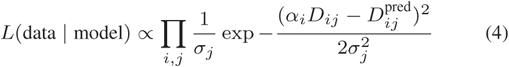

Here, *α*_*j*_ a are normalization values for each data condition *i*. The second term in Equation (3) is the prior, *P*(model):

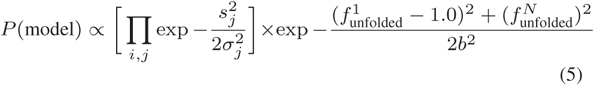

This prior prevents the estimated errors at each residue *σ*_*j*_ from dropping to zero; the *s*_*j*_ values set floors on *σ*_*j*_, and are chosen by default to be 10% of the mean data value at the residue *j*. The remaining terms in the prior ensure that the population fraction of the first state (unfolded) transitions from near 1.0 to near 0.0 (i.e., the experiment has been designed to capture the entire transition); the parameter *b* is chosen by default to be 0.05. All parameter choices can be changed by the user.

To visualize the full behavior of the posterior probability, the `LIFFT. m` function accepts a grid of candidate values for the input parameters (e.g., *K*_*d*_ and *n*). For each such setting of parameters, the tool optimizes the state chemical mapping patterns 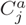 the fitted error at each residue *σ*_*j*_, and normalization values *α*_*i*_ (subject to the constraint 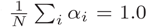) through iterative optimization (Supplemental Information). A refinement of the best parameter values is also carried out through MATLAB’s `fminsearch` functionality. Uncertainties in the fit parameters are estimated based on how far the parameters must be shifted from this best fit to observe the log-posterior drop by 2.0 (corresponding to two standard errors in each parameter if the uncertainties are Gaussian); these numbers provide convenient summary statistics for publication. The matrix of log posterior probability vs. each parameter setting is also returned for use in downstream statistical calculations; see, e.g., Supporting Information of (Frederiksen *et al.,* 2012). For a data set mapping 60 residues at 16 conditions and a 20 by 20 grid of input parameters, the calculation takes less than 2 seconds on a MacBook laptop (2.9 GHz Intel Core i7) with four CPU cores.

Figure 1 shows representative output for a Mg^2+^ binding curve measured for a domain of the signal recognition particle stabilized by two Rosetta-designed mutations, assessed by dimethyl sulfate probing (Das *et al.,* 2010)). In this case, the log-posterior plot (Fig. 1A) shows contours shaped like a tilted egg near the best fit: there are weak, non-linear correlations between uncertainties in the dissociation constant and apparent Hill coefficient. The fit is excellent, with low residuals (Fig. 1B), and gives 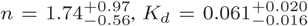. The final fits pass through the chemical reactivity data at user-selected residues that lie inside the SRP loop (Fig. 1C). A similar LIFFT quantitation for data for the wild type RNA 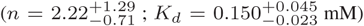 showed that the mutant required less Mg^2+^ to fold (confirming a stabilization by 1.1 ± 0.4 kcal/mol, using the thermodynamic framework described in (Lipfert *et al*., 2014); uncertainties are 2 standard errors).

**Fig. 1.**
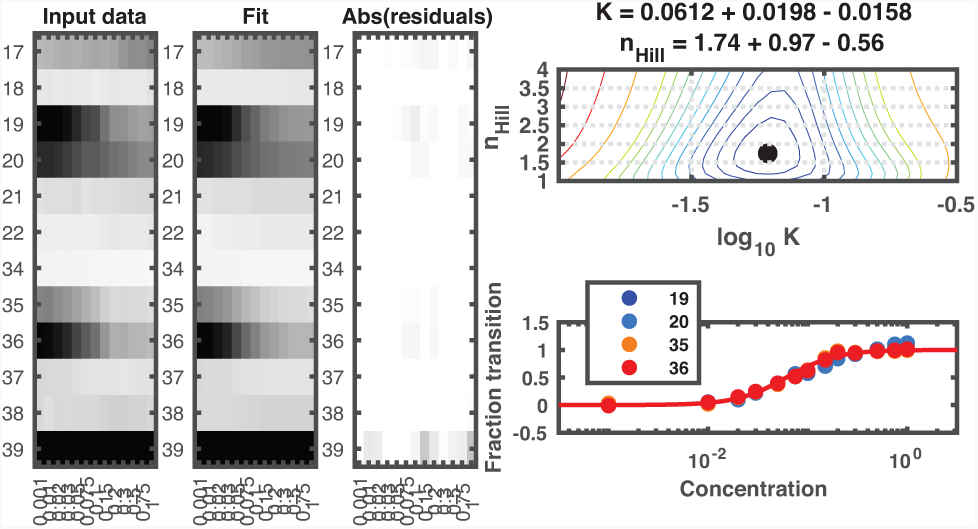
Output from LIFFT for Mg^2^ +-induced folding of a re-designed signal recognition particle RNA domain, probed by dimethyl suldate mapping. (A) Log-posterior contour map. (B) Heat maps of input data, best fit, and absolute values of residuals. (C) Fit curves overlaid on user-specified residues. Concentration units are mM. Data from (Das et al., 2010)

In addition to the Hill fit, several other thermodynamic models are available in LIFFT. For riboswitches that bind a single ligand, users can enforce *n* = 1 in the Hill fit, equation (2). For modeling molecules that bind up to two ligands, the Hill fit is incorrect (Kuriyan *et al.,* 2012), but the correct model is supplied with LIFFT (Kladwang *et al.,* 2012; Frederiksen *et al.,* 2012). Thermal melts (in which molecular structure is monitored vs. temperature) can be fit to models with constant enthalpy and entropy differences between states (Vaidyanathan *et al.,* 2017). Any other model that depends on up to two thermodynamic parameters can be supplied. Models with multiple transitions can be handled by separating out data subsets that define distinct transitions (Frederiksen *et al.,* 2012). Demonstrations on actual data sets for each of the above test cases are provided under `lifft_demo. m` and run successfully with MATLAB R2014a, R2016a, and R2017b on Mac OS 10.12.6.

## Acknowledgements

The author thanks members of the Das laboratory for testing LIFFT.

